# Polymeric Lysosomal-Targeting Chimeras: Extracellular Targeted Protein Degradation Without Co-opting Lysosome-Targeting Receptors

**DOI:** 10.1101/2024.09.18.613672

**Authors:** Ryan Hung-Hsun Lu, Jithu Krishna, Yasin Alp, S. Thayumanavan

## Abstract

Extracellular targeted protein degradation (eTPD) is an emerging modality to regulate protein levels without genomic interruption. Current strategies co-opt lysosome-targeting receptors (LTRs) that are ubiquitously present in most cells, offering a high success rate of eTPD across cell types and tissues. Opening up the binding complementarity requirement from LTRs to any overexpressed cell surface receptor offers to endow eTPD platforms with new cellular targeting capabilities. Here, we report polymeric lysosome-targeting chimeras (PolyTACs), a polymer-antibody conjugate based platform for the targeted degradation of membrane-bound and soluble proteins without the need for involving LTRs. Mechanistic investigations suggest a non-classical uptake pathway that is attributed to the membrane tension caused by the multivalent interaction between the PolyTACs and the overexpressed functionalities on the cell surface. The utility of PolyTACs in eTPD has been demonstrated with three therapeutically relevant membrane proteins. Additionally, the same design principle has also been leveraged to bind and drag soluble extracellular proteins into the lysosome. The design and fabrication simplicity, non-reliance on LTRs, and tissue-targeting capabilities open up new avenues for eTPD in many disease-specific applications.

## Introduction

Potent downregulation of the activity of a specific protein along with diminution of its scaffolding functions, without the need for a well-defined active site binding pocket in the target protein, have propelled targeted protein degradation (TPD) as the more versatile alternate to drug design based on occupancy based inhibitors (*1*–*3*). Conceptually, most of these degraders are based on two ligand functionalities brought together by a linker, where one of the ligands binds to the target protein of interest and the other ligand binds to an effector protein that ultimately directs the former to a cellular degradation machinery (*4*). This molecular chimera-based approach of leveraging the cell’s own machinery to downregulate specific protein function started with TPD of intracellular proteins using the proteasome and autophagosome, where the effector proteins are based on E3 ligases and autophagosome receptors respectively. Considering that extracellular proteins constitute >25% of the human proteome and are implicated in many important human pathologies (*5*), more recently, this concept was elegantly extended to extracellular proteins using lysosome targeting chimeras (LYTACs) (*6*–*8*). Here, the protein of interest is targeted by an antibody and a ligand conjugated to the antibody targets an effector protein based on lysosome-targeting receptors (LTRs), such as the mannose-6-phosphate receptor or the asialoglycoprotein receptor (*7*). While the advantage of leveraging the LTRs is that these are ubiquitously present in most cells, we posited that an LTR-independent extracellular TPD (eTPD) would offer new opportunities for degrading membrane proteins in specific cells or tissues. In this manuscript, we disclose a versatile LTR-independent polymeric lysosome targeting chimera (PolyTAC) platform with the potential for opening up new avenues for eTPD.

PolyTACs are based on antibody-polymer conjugates, where the antibody targets the specific extracellular protein for degradation, and the polymer is designed to make multivalent contact with the cell membrane. In designing these PolyTACs, we were inspired by a combination of three now well-accepted observations. First, targeted therapy is often based on overexpressed receptors on the surface of pathological cells and tissues (*9*). Therefore, incorporating multiple ligand moieties on a scaffold will enhance target specificity. Second, multivalency from ligand-bearing artificial scaffolds not only offers superselectivity to target the overexpressed receptors on cells but also enhances endocytosis-mediated cellular uptake (*10*–*13*). This suggests that if an antibody is conjugated to a polymer that makes a multivalent contact with a cell surface, it will cause both the receptor targeted by the antibody to be endocytosed along with the polymer. Third, the biggest challenge in delivering biologics to cells involves the difficulty in endosomal escape (*14, 15*), which means most of these scaffolds that are taken up through the endosome fuse with the lysosome. Put together, the antibody in the PolyTAC should bind to a specific targeted receptor, and when the antibody-conjugated polymer makes a multivalent contact with the cell surface, the scaffold along with the targeted cell surface protein should be shepherded to the lysosome for degradation.

For the multivalent contact with the cell surface, we envisaged leveraging the thiol moieties resulting from the overexpression of redox-controlling proteins on cancer cells due to their aberrant redox status (*16, 17*). The reduced cysteines, also known as exofacial thiols, on these cell surfaces, are intriguing biomarkers of oxidative stress and have been utilized for cell-specific targeting (*18*–*23*). Here, we report a plug-and-degrade PolyTAC platform, where the conjugation of thiol-reactive polymeric binders to antibodies converts the latter from target-specific inhibitors into target-specific degraders (Fig. 1a). This lysosomal trafficking approach that circumvents the dependence on lysosome-targeting and lysosomal-sorting proteins represents a paradigm shift, opening new avenues for cell-specific eTPD.

**Fig. 1.**
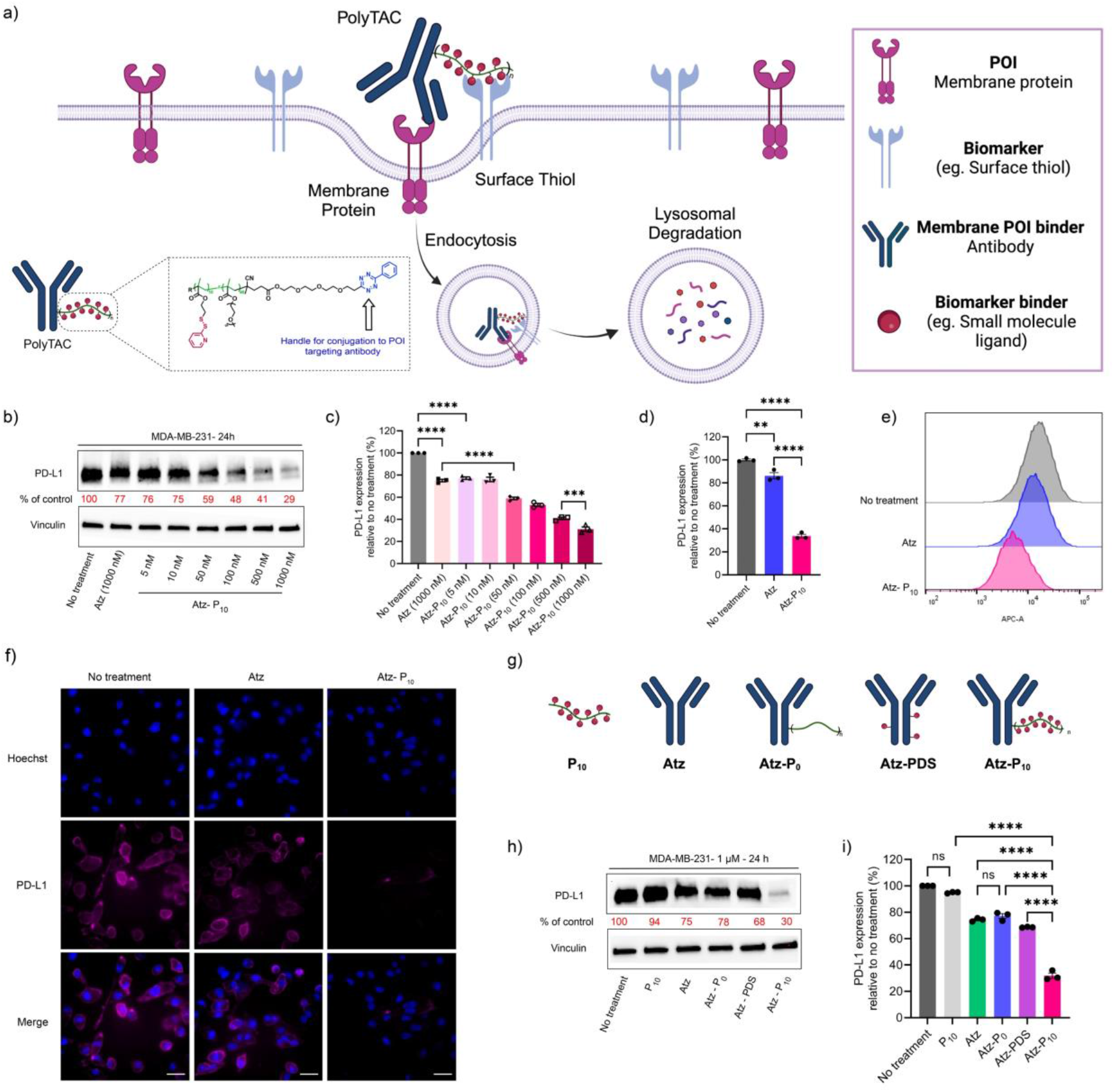
PD-L1 degradation mediated by Atz-P_10_ PolyTAC: **a**, Schematic representation of plug-and-degrade PolyTAC platform. **b-c**, Western blot analysis of total PD-L1 levels in MDA-MB-231 cells with the treatment of Atz at 1 µM or varying concentrations of Atz-P_10_ PolyTAC for 24 h. *n* = 3 biologically independent experiments. **d-e**, Relative surface expression of PD-L1 determined by live-cell flow cytometry following the treatment of 1 µM of Atz, Atz-P_10_ for 24 h. **f**, Visualization of PD-L1 degradation in MDA-MB-231 cells by confocal microscopy with the treatment of Atz or Atz-P_10_ at 1 µM for 24 h. Scale bar: 20 µm. **g**, schematic representation of PDS-containing polymers (P_10_), antibody (Atz), antibody with appended polymer with no PDS units (Atz-P_0_), antibody with one PDS unit per linker (Atz-PDS), and Atz-P_10_. **h-i**, Western blot analysis of total PD-L1 levels in MDA-MB-231 cells after the treatment with 1 µM P_10_, Atz, Atz-P_0_, Atz-PDS, and Atz-P_10_ for 24 h, suggesting the multivalent interactions behind PolyTACs for extracellular targeted protein degradation. *n* = 3 biologically independent experiments. The statistical significance was assessed using two-way ANOVA, *P < 0.05, **P < 0.01, ***P < 0.001, ****P < 0.0001.

## Results

### Construction of PolyTACs

To test the PolyTACs concept, we targeted the degradation of programmed death-ligand 1 (PD-L1), as it is also known to be overexpressed on the surface of many cancer cells (*24*). To test the possibility of leveraging thiol overexpression to degrade PD-L1, atezolizumab (Atz) and pyridyl disulfide (PDS) were used as the anti-PD-L1 antibody and thiol-reactive functionality in the polymer chain respectively in the PolyTAC, Atz-P_10_ (Fig. 1a). Reversible addition-fragmentation chain transfer polymerization was used to prepare the P_10_ methacrylate copolymer, containing ∼10 repeat units of the thiol-reactive PDS-ethyl methacrylate and ∼40 repeating units of the hydrophilic pentaethyleneglycol methacrylate (Fig. S1). A tetrazine unit was installed at the polymer chain terminus as the handle for conjugation with the antibody. The Atz antibody was modified with a trans-cyclooctene (TCO) functionality using an NHS-ester reaction with the lysines on the antibody (Fig. S2a) (*25*). PolyTACs were then successfully prepared *via* the click reaction between the tetrazine and the TCO moieties, as indicated by the migration of Atz-P_10_ and Atz-P_0_ away from Atz in gel electrophoresis (Fig. S2b). A combination of BCA assay and the relative fluorescence from Cy5-labeled P_10_ was used to characterize the Atz-P_10_ PolyTAC that contains an average of 3 polymer chains per antibody (Fig. S3).

### Degradation of PD-L1 using PolyTACs

We then evaluated PD-L1 degradation by Atz-P_10_ in MDA-MB-231 cells. This cell line is chosen for its overexpression of both PD-L1 and exofacial thiols. To further validate this, these cells were assessed by maleimide-R phycoerythrin (Mal-PE) staining and the observed high expression level of exofacial thiols (Fig S4) is consistent with prior studies (*16, 22, 23*). Similarly, PD-L1 expression was assessed by western blot. When these cells were treated with Atz-P_10_, a dose-dependent PD-L1 degradation was observed with a half-maximal degradation concentration (*DC*_50_) of ∼100 nM and a degradation maximum (*D*_max_) of ∼70% at 1 µM in 24 h (Fig. 1b, c). Higher potency is observed for 48 h treatment with a *DC*_50_ of 10 nM and a similar *D*_max_ (Fig. S5). Interestingly, the *D*_max_ is quite time-independent at least from 1 to 48 h. That is, at 1 µM concentration, PolyTACs show comparable degradation of PD-L1 (60-70%) in MDA-MB-231 cells for 1, 6, 12, 24, 48 h treatments (Fig. S6). These results suggest facile protein binding, trafficking, and degradation. With flow cytometry to probe the PD-L1 remnants on the cell surface, we observed a similar decrease in the surface expression of PD-L1 by ∼70% for Atz-P_10_ relative to no treatment in 24 h, while only ∼20% reduction was observed for Atz alone at the same concentration (1 µM) (Fig. 1d, e). Confocal laser scanning microscopy (CLSM) images also revealed a significant decrease in PD-L1 expression at the cell surface upon treatment with Atz-P_10_ compared to the treatment with Atz alone or no treatment (Fig. 1f), further supporting the observations in flow cytometry and western blot.

To test whether multivalency is required for PD-L1 degradation, the PD-L1 degradation capability of Atz-P_10_ was compared with several analogues as controls - including antibody alone (Atz), antibody with one PDS unit per linker (Atz-PDS), and antibody with appended polymer with no PDS units (Atz-P_0_). All controls exhibit a much lower ability to degrade PD-L1 even at 1 µM concentration, compared to Atz-P_10_ (Fig. 1g, h). Also, P_10_ alone exhibited an insignificant difference in PD-L1 levels, compared to the no treatment control. Moreover, co-treatment with individual P_10_ and Atz showed similar degradation efficiency to Atz alone, as did the subsequent treatment of Atz followed by P_10_ (Fig S7). These findings indicate that the antibody-polymer conjugates, Atz-P_10_, is an essential component to induce considerable PD-L1 degradation, suggesting the multivalent interactions as the driver behind the observed degradation with the PolyTAC.

### Mechanism of extracellular targeted protein degradation by PolyTACs

Next, we sought to understand the mechanism of PolyTAC-mediated degradation. To determine whether PolyTACs proceed with protein degradation via lysosomes or proteasomes, MDA-MB-231 cells were pre-treated with either PBS, bafilomycin A1 (Baf, an inhibitor of lysosome acidification), or MG132 (a proteasome inhibitor), followed by incubation with medium alone, Atz, or Atz-P_10_ (Fig. 2a, b). Baf dramatically compromised the PD-L1 degradation capability of Atz-P_10_, showing insignificant differences compared to Atz alone, whereas MG132 had no impact on PD-L1 degradation by Atz-P_10_. These findings suggest that PolyTACs mediate protein degradation by trafficking protein targets to lysosomes.

**Fig. 2.**
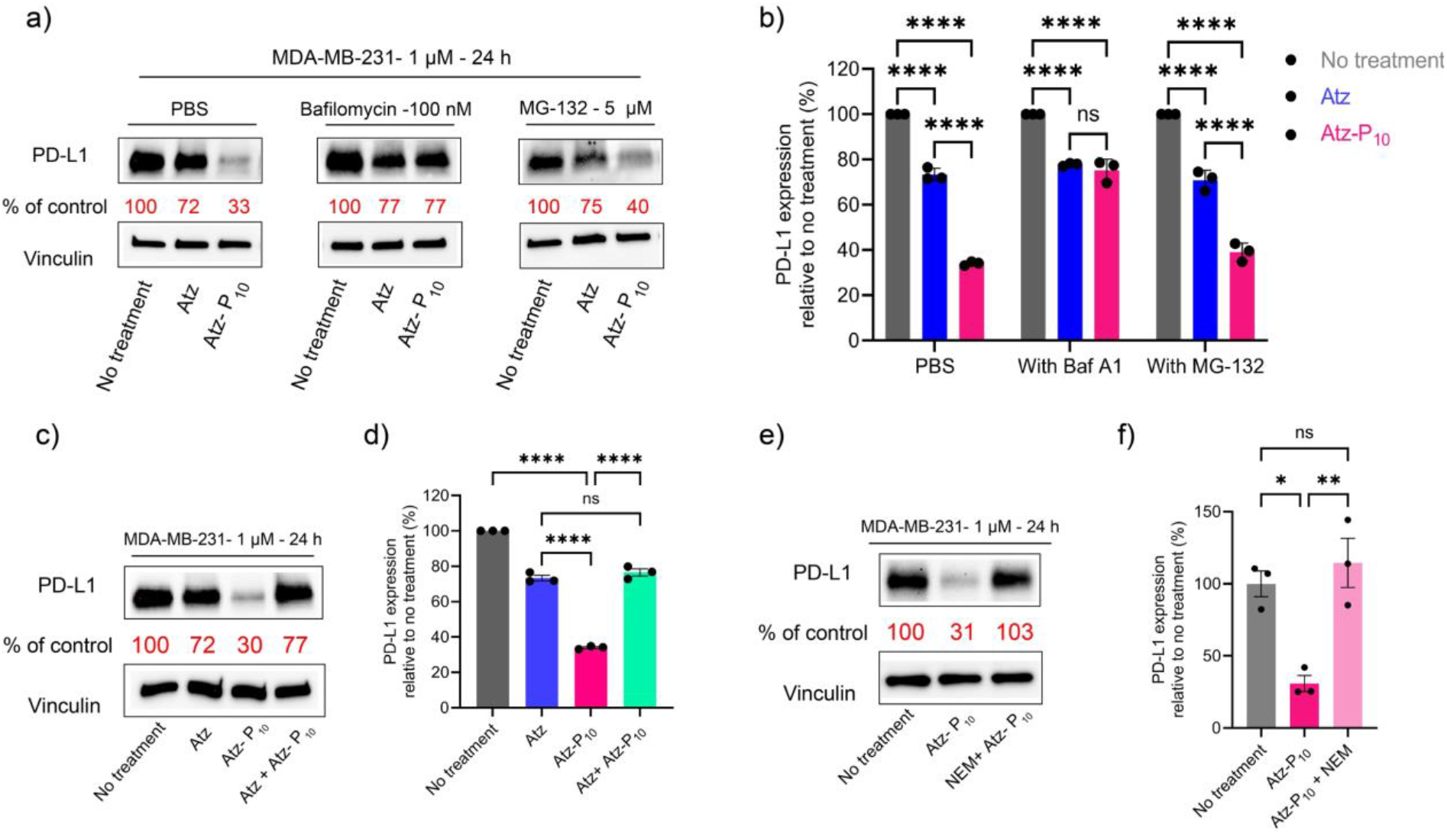
Mechanism of targeted protein degradation of membrane proteins by PolyTACs: **a-b**, Western blot analysis of total PD-L1 levels in MDA-MB-231 cells after the pretreatment for 1 h with either 100 nM bafilomycin or 5 µM MG-132 followed by 24 h of treatment with 1 µM of Atz, Atz-P_10_ PolyTAC degrades PD-L1 in a lysosome-dependent manner. *n* = 3 biologically independent experiments. **c-d**, Western blot analysis of total PD-L1 levels in MDA-MB-231 cells after the pretreatment of 1 µM of Atz for 1 h in MDA-MB-231 cells followed by 24 h of treatment with 1 µM of Atz-P_10_ PolyTAC shows the decreased degradation due to the masking of surface PD-L1 by Atz. *n* = 3 biologically independent experiments. **e-f**, MDA-MB-231 cells pre-treated with the thiol inhibitor *N*-ethylmaleimide (NEM) blocked the PD-L1 degradation capability of Atz-P_10_ showing the thiol mediated degradation of PolyTAC. *n* = 3 biologically independent experiments. The statistical significance was assessed using two-way ANOVA, *P < 0.05, **P < 0.01, ***P < 0.001, ****P < 0.0001.

Comparison of Atz-P_10_ with other controls in PD-L1 degradation suggests that the antibody binding to the protein target and the polymer-appended functionalities engaging with exofacial thiols are both critical factors for multivalency-induced endocytosis and subsequent degradation in the lysosome. First, we investigated whether the target protein binding by the antibody is essential for degradation. We evaluated the PD-L1 degradation in MDA-MB-231 cells when the cells were pre-treated with Atz for 1 h and then with Atz-P_10_ for 24 hours (Fig. 2c, d). The prior presence of Atz substantially diminished the protein degradation capability of Atz-P_10_, showing similar PD-L1 degradation to that of Atz alone. This is attributed to the masking of cell surface PD-L1 by Atz and therefore preventing Atz-P_10_ from engaging with PD-L1.

Next, we explored the role of exofacial thiols in the PolyTAC. *N*-ethylmaleimide (NEM) has been shown to be effective in blocking the exofacial thiols via thiol-maleimide conjugation to inhibit the thiol-mediated uptake (*22, 23*). If the thiol-PDS interaction is the key to the observed multivalency-driven PD-L1 degradation, then pre-treating MDA-MB-231 cells with NEM should impair the degradation capability of Atz-P_10_. Indeed, the degradation ability of Atz-P_10_ is completely muted when pre-treated with NEM (Fig. 2e, f). Together, these findings indicate that both antibody and the polymer-appended PDS functionalities are critical to initiating endocytosis and lysosomal trafficking for effective eTPD.

### Cellular internalization of PolyTACs

We further investigated the endocytic mechanisms by which PolyTACs cause the lysosomal degradation of membrane proteins. To explore potential endocytic mechanisms, we studied PD-L1 degradation mediated by Atz-P_10_ in the presence of various inhibitors (Fig. S8a). Inhibitors of clathrin-mediated endocytosis (dynasore, pitstop 2, and chlorpromazine) did not significantly impact PD-L1 degradation compared to controls. Similarly, inhibitors of caveolae-mediated endocytosis (methyl-*β*-cyclodextrin and genistein) and an inhibitor of macropinocytosis (5-(*N*-ethyl-*N*-isopropyl)-amiloride) had negligible effects on the degradation capability of Atz-P_10_. These results suggest that PolyTACs induce endocytosis and lysosomal trafficking via clathrin- and caveolae-independent pathways.

We observed that inhibition of endolysosomal acidification by bafilomycin completely compromises PD-L1 degradation (Fig. 2a), which suggests an endosomal pathway for PolyTAC entry into cells. Similarly, pre-blocking the protein target or the exofacial thiols also suppresses PolyTAC-mediated protein degradation (Fig. 2c, e), which reiterates the crucial role of multivalent interactions. With these observations, combined with the lack of dependence on the classical endocytosis inhibitors, we propose that the endocytic uptake in the case of PolyTACs is likely caused by the polyvalent interactions induced membrane tension. Polyvalent interactions between the polymer and the cell membrane associated thiols concurrently offer both adhesion energy and membrane deformation to cause atypical endocytosis. In fact, there is some literature precedence for this possibility. Lipid-binding virions and virus-like materials have been shown to induce membrane deformation and clathrin-independent endocytosis through multivalent lipid binding (*26*). Similarly, the engulfment of giant unilamellar vesicles, mediated by multivalent interactions with nanoparticles, has been shown to be driven by multivalency-induced membrane tension (*27*). If the multivalency-based adhesion energy indeed offers the driving force for the endocytic uptake, we hypothesized that we should observe protein degradation by PolyTACs even at 4 °C. Note that classical endocytosis pathways are significantly muted at 4 °C. While the observed degradation of PD-L1 by Atz-P_10_ was about ∼59% at 37 °C for 1 h, the degree of degradation decreased only to ∼39% at 4 °C (Fig. S8b). Taken together, these findings suggest that multivalency through antibody/surface protein binding and PDS/exofacial thiol interactions elevates cell membrane tension and promotes endosome formation, ultimately trafficking antibody-bound protein targets into the lysosome for degradation.

### Testing the PolyTACs scope with other cell surface proteins

To investigate the broad applicability of this platform for the degradation of different surface protein targets with overexpressed exofacial thiols, we also assembled PolyTACs of Sac-P_10_ and Cet-P_10_ that are based on sacituzumab and cetuximab antibodies respectively (Fig. 3a). These antibodies bind to trophoblast cell surface antigen 2 (Trop2) and epidermal growth factor receptor (EGFR), overexpressed in SKBR3 and MDA-MB-231 cells respectively.

**Fig. 3.**
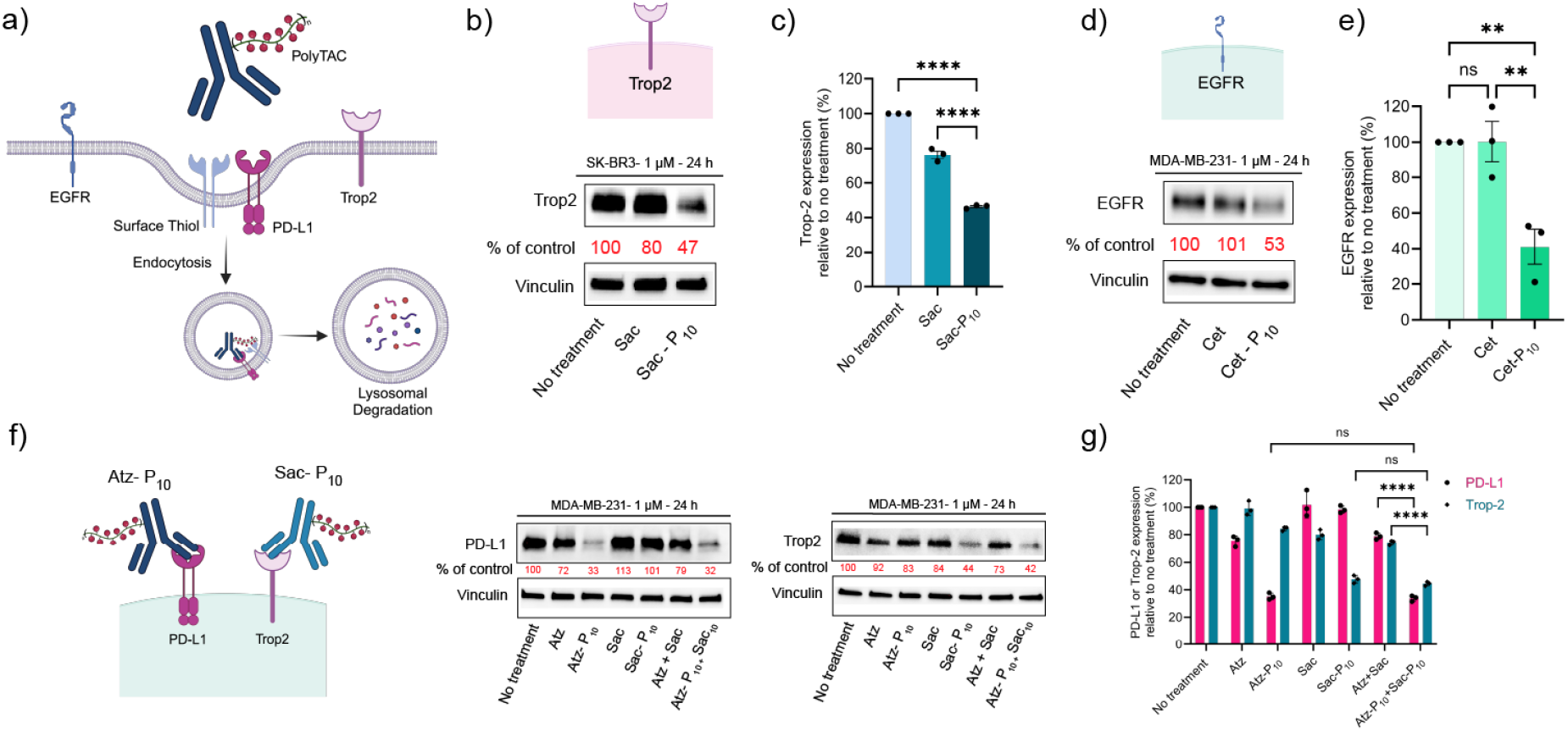
Degradation of other therapeutically relevant cell surface proteins by PolyTACs: **a**, Schematic of the PolyTACs for targeting different cell surface proteins for lysosomal degradation. **b-c**, Trop2 levels were significantly reduced after a 24 h treatment of SK-BR3 cells with 1 µM of Sac-P_10_ PolyTAC compared to Sac alone. n = 3 biologically independent experiments. **d-e**, EGFR levels were reduced to half after a 48 h treatment of MDA-MB-231 cells with 1 µM of Cet-P_10_ PolyTAC compared to Cet alone. n = 3 biologically independent experiments. **f-g**, The consecutive degradation of PD-L1 and Trop2 in MDA-MB-231 cells after the treatment of 1 µM of both Atz-P_10_ PolyTAC and Sac-P_10_ PolyTAC for 24 h. n = 3 biologically independent experiments. The statistical significance was assessed using two-way ANOVA, *P < 0.05, **P < 0.01, ***P < 0.001, ****P < 0.0001.

Trop2 is a type-I transmembrane glycoprotein that is overexpressed in a wide range of epithelial cancers (*28*) and has been a target for oncology, highlighted by the recent approval of an antibody-drug conjugate (ADC), Trodelvy, for triple negative breast cancer (*29*). The P_10_ polymer was conjugated on to sacituzumab using the procedures described above for Atz-P_10_ to produce the Sac-P_10_. Treatment of SKBR3 cells with 1 µM concentration of this PolyTAC caused a decrease in Trop2 levels by ∼55%, while the sacituzumab antibody by itself caused only ∼25% reduction, as quantified by western blot (Fig. 3b, c). As Trop2 is also overexpressed in MDA-MB-231 cells, we also tested the effect of Sac-P_10_ in these cells and found that Sac-P_10_ causes a substantially better degradation of Trop2 (∼65%) too, compared to the antibody alone (∼10%). (Fig. S9) The cell-line dependent degree of degradation of specific epitopes is attributed to the inherent internalization efficiency of cells, abundance of the cell surface protein target, and expression level of exofacial thiols (*30, 31*).

Next, we targeted EGFR, a transmembrane glycoprotein that is also upregulated in many cancer cells, along with the exofacial thiols (*32*). We incorporated the P_10_ with cetuximab (Cet), an FDA-approved anti-EGFR antibody, to achieve the Cet-P_10_ for EGFR degradation. The treatment of Cet-P_10_ at 1 µM for 48 h in MDA-MB-231 cells caused 60% EGFR degradation, whereas negligible degradation was observed in the treatment of Cet alone at the same concentration (Fig. 3d, e). We further tested the combination degradation using PolyTACs targeting different surface proteins (Fig. 3f). MDA-MB-231 cells were co-treated with Atz-P_10_ and Sac-P_10_ for 24 hours, followed by the quantification of remaining PD-L1 and Trop-2 expression. The co-treatment demonstrated comparable degradation capability of PD-L1 and Trop-2 to that of using individual degraders (Fig. 3f,g), indicating that sufficient exofacial thiols are present to initiate concurrent degradation of two different proteins.

### PolyTACs enable intracellular uptake of soluble extracellular proteins

Next, we tested the possibility of the antibody component of the PolyTACs binding to soluble extracellular proteins and leveraging the polyvalent thiol-disulfide interaction to transport them into cells and traffic them to the lysosome (Fig. 4a). Soluble proteins, such as vascular endothelial growth factor (VEGF) and Immunoglobulin G (IgG), are associated with signal transductions in cancer and autoimmune diseases (*33*). Therefore, there is an interest in recruiting these soluble extracellular proteins inside the cells and trafficking them to the lysosome. As a proof-of-concept, we first targeted rabbit-IgG to determine whether PolyTACs could drag soluble proteins with different constructs into cells via multivalency. The P_10_ polymer was conjugated to goat anti-rabbit IgG (α-rabbit IgG) to produce PolyTACs (α-rabbit IgG-P_10_), with ∼3 polymer chains per antibody. We first investigated whether the α-rabbit IgG-P_10_ would promote cellular uptake of AF647-labeled rabbit IgG in comparison to α-rabbit IgG alone. From flow cytometry, a 5-fold increase in MFI of AF647-labeled rabbit IgG was observed in the treatment that contains α-rabbit IgG-P_10_, relative to rabbit IgG alone. Also, there is no difference in the uptake of rabbit IgG when the treatment contains α-rabbit IgG, relative to rabbit IgG alone (Fig. 4b). This once again confirms the polyvalency role of the polymer and the antibody binding to the target soluble protein for trafficking. This assertion is further supported by the substantially enhanced signal of AF647 from rabbit IgG plus α-rabbit IgG-P_10_ in CLSM images (Fig. 4e). The AF647 signal from rabbit IgG plus α-rabbit IgG-P_10_ was mainly co-localized with LysoTracker Green (Fig. S10), demonstrating the rabbit IgG mainly trapped inside the lysosome following endocytosis.

**Fig. 4.**
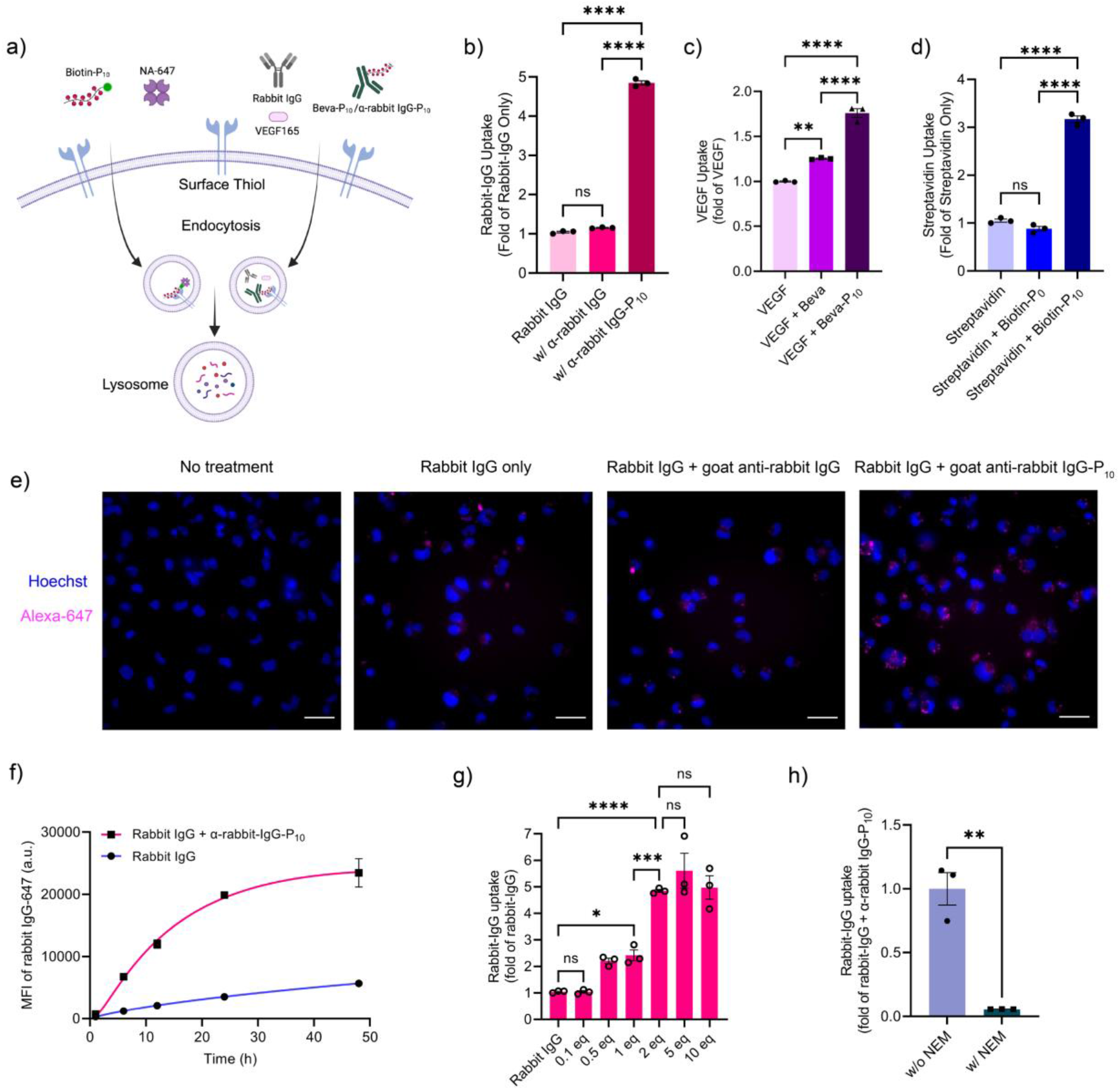
PolyTACs enable intracellular uptake of soluble extracellular proteins: **a**, Schematic of PolyTAC concept for targeting soluble extracellular proteins for lysosomal degradation. **b**, Flow cytometry results showing the enhanced uptake of rabbit IgG when treated with Goat anti-rabbit IgG-P_10_ for 6 h in MDA-MB-231 cells, Rabbit IgG alone shows similar uptake as that with the treatment of Goat anti-rabbit IgG. n = 3 biologically independent experiments. **c**, flow cytometry bar graphs showing significant uptake of VEGF–647 in MDA-MB-231 cells following 6 h of treatment with Beva-P_10_ compared to treatment with Beva alone and VEGF alone. Beva control showed no difference compared to treatment with VEGF–647 alone. n = 3 biologically independent experiments. **d**, Flow cytometry results demonstrating a significant uptake of Streptavidin (NA-647) in MDA-MB-231 cells treated for 6 h with Biotin-P_10_ compared with NA-647 alone or NA-647+ Biotin-P_0_. n = 3 biologically independent experiments. **e**, CLSM images showing the enhanced uptake of rabbit IgG when treated with Goat anti-rabbit IgG-P_10_ for 6 h in MDA-MB-231 cells compared to the treatment with either Rabbit IgG alone or Rabbit IgG with Goat anti-rabbit IgG. n = 3 biologically independent experiments. Scale bar; 20 µm. **f**, Time-dependent IgG uptake using flow cytometry shows the highest increment in cellular internalization relative to IgG alone by 6 folds in 6 h and steady state in 48 h. n = 3 biologically independent experiments. **g**, the flow cytometry data shows that the levels of IgG uptake depend on the equivalence of PolyTAC. IgG uptake steadily increases from 0.1 equivalence of α-rabbit IgG-P_10_ and reaches a plateau at 2 equivalences. n = 3 biologically independent experiments. **h**, Pre-blocking the exofacial thiols of MDA-MB-231 cells with the thiol inhibitor NEM resulted in a fivefold decrease in IgG uptake with Goat anti-rabbit IgG-P_10_ compared to the control without NEM. n = 3 biologically independent experiments. The statistical significance was assessed using two-way ANOVA, *P < 0.05, **P < 0.01, ***P < 0.001, ****P < 0.0001.

We further tested the uptake and trafficking using a smaller soluble protein VEGF by conjugating bevacizumab (Beva), an antibody against VEGF, to P_10_ to generate the Beva-P_10_. MDA-MB-231 cells were incubated with FITC-labeled VEGF165 (FITC-VEGF) along with Beva or Beva-P_10_. We observed an1.8-fold increase in MFI of FITC-VEGF in the presence of Beva-P_10_, but only a 1.2-fold increase in the presence of Beva, relative to the FITC-VEGF by itself (Fig. 4c). Beva itself is known to possess some propensity to be taken up by the cells (*34*). The increase in VEGF uptake in the presence of Beva-P_10_ further supports the importance of multivalent interactions between exofacial thiols and PolyTACs.

Next, we wanted to extend this idea to a small molecule ligand instead of the antibody in the PolyTAC. NeutrAvidin-647 (NA-647), an Alexa Fluor-647 (AF647)-labeled protein, was used as a model protein to test for PolyTAC-driven protein uptake and lysosomal trafficking. Accordingly, biotinylated-P_10_ (Biotin-P_10_) and biotinylated-P_0_ (Biotin-P_0_) were synthesized. Biotin-P_10_ was synthesized by first obtaining a biotinylated RAFT reagent, which was then co-polymerized with PDS-methacrylate and PEG-methacrylate. The same RAFT reagent was used to polymerize PEG-methacrylate alone to generate Biotin-P_0_. (Fig. S11). When incubated with NA-647 alone or NA-647 plus Biotin-P_10_ or Biotin-P_0_ for 24 h, Biotin-P_10_ clearly exhibits increased fluorescence in cells by 3-fold compared with NA-647 alone, whereas Biotin-P_0_ showed a negligible difference in fluorescence relative to NA-647 alone (Fig. 4d). This flow cytometry based results were further supported by fluorescence microscopy experiments based on CLSM (Fig. S12). High colocalization of green and red fluorescence, from lysotracker and NA-647 respectively, indicates that Biotin-P_10_ successfully traffics the complex of NA-647 plus Biotin-P_10_ into lysosome. These results, together with the observed PolyTAC-mediated cellular uptake and trafficking of α-rabbit IgG and VEGF, show that PolyTACs can serve to combine the receptor-ligand interactions with a soluble extracellular protein with multivalent interaction between exofacial thiols to promote cellular uptake and lysosomal trafficking.

Next, we evaluated the temporal evolution of the PolyTAC-mediated uptake with α-rabbit IgG as the target protein. Cellular uptake of IgG occurred in a time-dependent manner with the highest increment in cellular internalization, relative to IgG alone by ∼5 folds in 6 h. A steady increase in the uptake is observed with time that appears to saturate around 48 h (Fig. 4f). Comparison of the initial rate for the PolyTAC uptake with the rate observed for the Rabbit-IgG by itself translates to ∼5.5 times higher rate of cellular uptake by the PolyTAC. We also evaluated the impact of the amount of PolyTAC relative to that of the target Rabbit IgG in the medium. At 0.1 equivalent of α-rabbit IgG-P_10_ PolyTAC, no observable difference is seen relative to the control. At 0.5 equivalent, a modest increase of ∼2 is observed. At 2 equivalents or more, a 5x enhancement is observed (Fig. 4g). Next, we evaluated whether the multivalency between the exofacial thiols and the disulfides in the PolyTACs is responsible for the observed enhancements in the soluble extracellular protein uptake. Indeed, when the exofacial thiols of the cells were pre-blocked by NEM, the observed increase in IgG uptake was completely nullified (Fig. 4h).

To ascertain whether the PolyTAC uptake mechanism along with the soluble extracellular proteins is similar to that observed for the membrane-bound proteins above, we first evaluated the uptake efficiency of rabbit IgG in the presence of the corresponding PolyTAC α-rabbit IgG-P_10_ at 4 ºC (Fig. S13a). Compared to the same experiment at 37 ºC, the IgG level in MDA-MB-231 cells at 4 ºC was reduced by 95%, as determined by flow cytometry. The uptake experiment at 4 ºC significantly affects endocytosis by inhibiting the process. This suggests that PolyTAC-mediated uptake is energy-dependent. While the degradation of membrane-bound proteins is not drastically affected at 4 ºC, the impact of lower temperature on the uptake efficiency of soluble proteins is significant. Next, we investigated the endocytic pathways by evaluating the rabbit IgG uptake efficiency in the presence of the PolyTAC, when pretreated with various inhibitors for clathrin-mediated endocytosis, caveolae-mediated endocytosis, and macropinocytosis (Fig. S13b-c). Similar to that observed with membrane-bound proteins, none of the inhibitors significantly reduced the PolyTAC-mediated uptake of rabbit IgG. These results suggest that the uptake pathway of PolyTACs for both soluble extracellular proteins and membrane-bound proteins is likely driven by the polyvalency-based cell membrane tension. The modest temperature dependence in the case of membrane protein uptake and a stronger dependence in the case of soluble extracellular proteins suggests the strong adhesion offered by the high binding affinity of antibody with its cell surface epitope, in addition to the multivalent interaction from the polymer, offers a significant driving force for the membrane deformation based cellular uptake in the former case. Overall, our findings indicate that multivalency is the major driving force directing both soluble and membrane-bound proteins into the intracellular domain *via* a membrane stress-driven endocytosis that ultimately leads to lysosomal degradation of the target proteins.

## Conclusion

In summary, our studies show that PolyTACs is a versatile platform for extracellular targeted degradation for both membrane-bound and soluble proteins. We find that PolyTAC-mediated protein degradation is driven by the binding of antibodies to protein targets, combined with the multivalent interaction between specific functionalities on the polymer and on the cell surface. These simpler requirements obviate the need to co-opt LTRs that open up opportunities for cell-specific targeting, such as in cells that display a high density of exofacial thiols. Evaluating the cellular uptake and degradation capability of PolyTACs in the presence of various cellular pathway inhibitors indicates that the observed eTPD indeed occurs through lysosomal degradation. Interestingly, we also find that cellular uptake does not occur through classical pathways such as micropinocytosis, clathrin-mediated, or caveolae-mediated endocytosis. We provisionally propose that the process is driven by the membrane stress generated by the multivalent interactions between the PolyTAC and the complementary cell surface functionalities. This LTR-independent access to eTPD introduces an arguably nimble molecular design paradigm that opens up new opportunities in many diseases, especially in the context of cell-or tissue-specific protein degradation. Because when the LTR-dependence is relaxed, the ligands on the polymer chain can now be used for any cell surface receptors, as demonstrated with exofacial thiols here. A simple, yet critical, feature of these PolyTACs is that these can be assembled rapidly with great synthetic ease, a key element in the possibility of ultimately translating these findings to the clinic. Overall, we anticipate that the PolyTAC platform will offer a general strategy for eTPD with an impact in therapies beyond what is currently possible.

## Acknowledgment

We acknowledge National Institutes of Health (GM-136395 and T32 GM135096) for the financial support. We thank Dr. Amy Burnside at the flow cytometry core facility for the training and discussions. We thank the UMass Amherst Institute of Applied Life Sciences (IALS) light microscopy facility (RRID: SCR_021148) and staff Dr. James Chambers for the guidance. Some of the Figures were created with BioRender.com.

